# Modelling phenotypic plasticity in cancer invasion and metastasis: from microscopic interactions to macroscopic dynamics

**DOI:** 10.64898/2026.07.17.739105

**Authors:** Dimitrios Katsaounis, Mark A.J. Chaplain, Nikolaos Sfakianakis

## Abstract

Cancer progression is driven by the interplay between cancer cell phenotypic variability regulated by EMT/MET, and cell-cell and cell-matrix interactions. Starting with an individual-based model, in which every cell is characterised by its position, velocity and a continuously varying epithelial-mesenchymal phenotype, we derive, via a kinetic description and a mean-field limit, two alternative macroscopic formulations: an Euler-like system that couples mass and momentum to the phenotypic variable, and a single advection-aggregation-diffusion equation (AADE) for the cancer cell density. Both macroscopic models retain the non-local adhesion-repulsion forces, haptotactic response to an evolving ECM, and phenotype-dependent transition dynamics driven by TGF-β. Numerical experiments in two spatial dimensions indicate that the macroscopic equations reproduce key scenarios obtained at the individual scale. In particular, a microscopic-macroscopic comparison shows that the AADE density reproduces acurately both the spatial localisation and the phenotypic decomposition of the individual cell population. We also demonstrate that varying only the steepness of the TGF-β switch function, changes the EMT response from an almost binary epithelial-mesenchymal (EM) separation to a partial EM phenotypes. This study provides a systematic bridge from stochastic, heterogeneous cell dynamics to continuum descriptions, for investigating phenotype driven tumour invasion and supporting the choice of macroscopic models in large-scale simulations and analytical studies.

## 1 Introduction

Cancer cell heterogeneity is a fundamental aspect of cancer development that influences tumour growth and response to treatment. Accordingly, the mathematical modelling of cancer development and therapies, necessitates the incorporation of the diverse phenotypic characteristics exhibited by the cancer cells. This diversity can give rise to complex dynamical phenomena, including varied interactions with the tumour microenvironment, competition between cell subpopulations, and the emergence of drug resistance. A key consequence of this phenotypic variability is the emergence of distinct cellular behaviours, in terms of motility, invasiveness, and interactions with the surrounding microenvironment.

Various mathematical modelling approaches have been proposed in the literature for the description of individual cancer cell movement. A number of these models are inspired from physical systems, particularly those describing interacting particle populations, where collisions between agents can generate complex dynamical patterns [27, 56, 20]. In these approaches, it is typically assumed that all agents possess identical physiological properties and exhibit homogeneous behavioural traits, both in terms of biased-random motion and interactions with their neighbours. While such assumptions are reasonable when modelling microparticles, e.g. molecules, they are not clearly appropriate in the case of biological entities, such as cells. In the case of cancer, in particular, it is observed that different phenotypic states are associated with distinct behaviours. For instance, certain phenotypes exhibit limited proliferative capacity while, on the other hand, they exhibit enhanced motility potential. Cancer cells may undergo phenotypic transitions between these states via processes such as epithelial-to-mesenchymal transition (EMT) and mesenchymal-to-epithelial transition (MET), resulting in a highly dynamic and diverse tumour cell population [52].

This paper aims to introduce a new individual-based model (IBM) of an interacting heterogeneous cancer cell population and to investigate the behaviour of macroscopic partial differential equation (PDE) models derived from the microscopic description, with a focus on capturing key migratory patterns. The derivation of such macroscopic models from individual-level dynamics has attracted considerable attention in various fields, particularly in the study of collective behaviour in animal populations, such as bird flocks and fish schools [56, 7]. The framework of kinetic theory, originally developed in the context of physical systems [17, 25, 48], provides the theoretical foundation for these derivations and has subsequently been adapted to biological settings.

Classical derivations within this framework typically assume that individuals share identical physiological characteristics, with heterogeneity arising only from spatial or stochastic effects. The derivation of the corresponding equations, which involves the passage from a finite number of individuals to the limit of an infinite population, is mathematically involved; see e.g. [7, 46, 3, 5, 4, 18]. Phenotype-dependent models for cancer growth and invasion have been considered before [44, 12, 15, 19, 23], where on-lattice agent-based models are considered and the corresponding macroscopic models are derived. Several other works have focused on the dynamic behaviour of phenotype-structured macroscopic PDEs alone [58, 14, 9]. In contrast, here we propose an off-lattice model, inspired by the framework introduced in [20], where cellular movement is described through Newton’s laws of motion and where phenotypic variability directly influences cell behaviour.

A further motivation behind this work is that continuum models should not merely reproduce the spatial distribution of cancer cells but also maintain phenotypic information from the individual cell level. This information includes cell migration and how this is regulated, response to environmental cues, and more. These all become particularly relevant for EMT, where the cell-population appears either with a clear, almost binary, distinction between the extreme epithelial-mesenchymal (EM) states or with a coexistence of various intermediate EM phenotypes. These two behaviours are also biologically very relevant: A steep response is consistent with systems exhibiting extreme EM phenotypes like breast cancer cell lines such as MCF7 or T47D, in contrast to the more mesenchymal-like cell lines such as MDA-MB-231 or SUM159 [1, 29]. A smoother response that allows intermediate EM phenotypes to coexist, appears in systems such as H1975 and PMC42-LA [59, 31, 30, 1]. This distinction is even more important, as partial EMT phenotypes have been associated with collective migration and increased metastatic capacity [47, 31, 30]. A schematic representation of the partial EMT and MET processes is shown in Figure 1

**Fig. 1:**
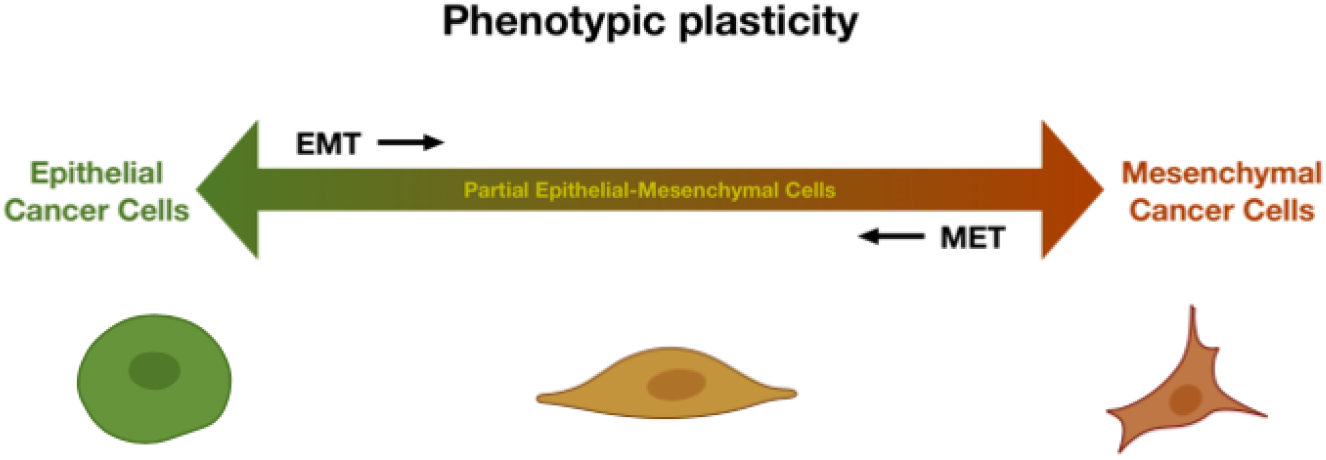
Schematic representation of the continuous phenotypic scale between epithelial and mesenchymal phenotypes where EMT and MET gives rise to partial epithelial-mesenchymal cells [33].

Motivated by these biological observations, we introduce an IBM that explicitly incorporates phenotypic dependence and subsequently we derive the corresponding macroscopic equations. The resulting framework allows us to investigate how phenotypic variability affects collective cancer cell dynamics while maintaining the connection between individual cell behaviour and the corresponding continuum description.

We then present numerical simulations to assess the behaviour of the derived models and their capacity to reproduce key features of cancer invasion dynamics. By comparing simulations of the individual cell dynamics with the corresponding advection-aggregation-diffusion equation (AADE), we show that the continuum density closely follows both the spatial support and the phenotypic composition of the underlying individual-based cell population. Moreover, changing only the steepness of the TGF-β-dependent switch function is sufficient to move the system from the endpoint-dominated EM regime to one in which partial EM phenotypes coexist. This establishes a link between cell phenotypic plasticity, continuum invasion models, and experimentally observed differences between binary and partial EMT behaviour.

The rest of the paper is structured as follows: in section 2 we present the related phenotype-dependent IBM, and then in section 3 we derive the corresponding macroscopic PDEs. Numerical simulations of both the microscopic and macroscopic models are presented in section 4, with the final section 5 providing a discussion of the results of the paper and outlining avenues for future work.

## 2 Modelling individual cancer cells

The diverse range of phenotypic characteristics that cancer cells exhibit plays a fundamental role in the progression of cancer. It is well established that, in particular, the primary cell population consists of epithelial-like cancer cells (ECCs). The ECCs possess the capability to undergo phenotypic transitions via the EMT process, leading to the generation of so called mesenchymal-like cancer cells (MCCs). The MCCs, characterised by enhanced migratory properties, facilitate, through the reverse MET, the formation of new micro-metastases distant from the primary tumour location. The heterogeneity of the tumour microenvironment highlights the complexity of the metastatic cascade, where the whole spectrum of cancer cell phenotypes contributes to the establishment of new tumour colonies and metastatic sites.

In this section, the objective is to describe these dynamics at the level of individual cancer cells, of a given phenotype, and their interactions with neighbouring cells and the surrounding microenvironment. In contrast to the more typical phenotypic classification into (binary) ECCs and MCCs as done in [50, 32], we consider here a continuous phenotypic spectrum; through partial EMT and MET processes, cancer cells of intermediate phenotype arise that display both epithelial and mesenchymal characteristics to varying degrees. The IBM considered here is extension of interacting-agent models such as [20, 56, 6], where the movement of the cell-agents follow Newton’s laws of motion. Let us note here that the proposed model is an off-lattice agent-based model, and differs from previous works considering on-lattice models and phenotype-dependent models, see e.g. [42].

We will begin by addressing the kinematics of cell migration, with particular attention to the interactions between cancer cells and their environment. It is assumed that cancer cells are able to sense and respond to external biochemical and biomechanical cues originating from the tumour microenvironment (TME). A key constituent of the TME is the extracellular matrix (ECM), which cancer cells utilise for navigation through the tissue. This is achieved via the formation of adhesion bonds with chemoattractants bound to collagen fibers in the ECM. The resulting phenomenon, called *haptotaxis*, describes the preferential migration of cancer cells towards regions of higher ECM density. In addition to haptotaxis, cancer cells interact with the environment through *chemotaxis*, where migration is directed by gradients of soluble biochemical substances, such as oxygen, glucose, or other nutrients, which are useful for cell survival. Both haptotaxis and chemotaxis contribute to the biased, directed motion of cancer cells towards regions of the TME that are favourable for tumour progression.

Another significant mechanism that we incorporate into the IBM accounts for cell-cell interactions. Specifically, we consider the physical space occupied by each cancer cell, assuming that cells cannot be arbitrarily close to each other. At very short distances, repulsion forces emerge between the cells, that prevent excessive proximity and induce mutual displacement. Furthermore, we include cell-cell adhesion mechanisms, where transmembrane glycoproteins called *cadherins* form biomechanical bonds between the membranes of interacting cells within a specific interaction range. These long-range interaction forces between neighbouring cancer cells can, in turn, promote the clustering of cancer cells and influence their collective dynamics.

In the current modelling framework, the equations that dictate the biological properties that affect cell movement are assumed to be:

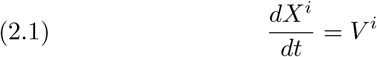

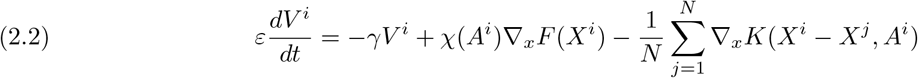

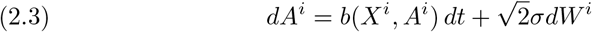

where *X*^*i*^ = *X*^*i*^(*t*) ∈ ℝ^*d*^, *V* ^*i*^ = *V* ^*i*^(*t*) ∈ ℝ^*d*^, *A*^*i*^ = *A*^*i*^(*t*) ∈ [0, 1] represent the position, velocity and the phenotype of the individual cell *i* = 1, …, *N* . Equation (2.1) describes the velocity, (2.2) the acceleration, and (2.3) the rate of change of the phenotype for each individual cell. In comparison to models such as the D’Orsogna model [20], here we have the addition of this free variable that describes the phenotypic changes of each individual cell.

The system (2.1)–(2.3) accounts for the modelling assumptions described above. In particular, the first term on the right-hand side of (2.2) represents a velocity damping effect, with a rate *γ >* 0, due to the physical friction between the cancer cells and the extracellular environment. The second term in (2.2) models the directed migration of cells towards regions of higher values of the function *F* : ℝ^*d*^ → ℝ that represents the density of the ECM or, in the case of chemotaxis, of the extracellular chemical signal. The corresponding rate of directed motion, denoted by *χ* : ℝ → ℝ, is assumed to depend on the phenotypic state *A*^*i*^ of the cell, thereby reflecting the heterogeneity of migratory behaviour among cancer cell populations. The final term in (2.2) describes the interactions among cancer cells through the interaction kernel *K* : ℝ^*d*^ → ℝ^*d*^. This kernel serves a dual purpose: modelling both the repulsive forces at short intercellular distances and the adhesive forces operative within an attraction range. Note that the adhesive interactions are assumed to depend on the phenotypic state *A*^*i*^ of the cells, and are consistent with the remark that mesenchymal-like cells exhibit reduced adhesion capabilities compared to the more epithelial ones. Finally, the factor *ε >* 0 accounts for the inertia effects of cellular movement.

The last equation of the system, (2.3), is formulated as a stochastic differential equation (SDE) that governs the phenotypic transitions of the cancer cells. The drift function *b* defines the deterministic component of the phenotypic dynamics, facilitating transitions between epithelial and mesenchymal states. The stochastic component (2.3) is characterised by a constant rate *σ >* 0, with *W* denoting a standard Wiener process. This stochastic term accounts for random fluctuations in the phenotypic state, which may be interpreted as modelling the occurrence of random mutations leading to phenotypic variation.

For the purposes of this work, and in the numerical simulations presented later, it is assumed that the phenotypic state variables *A*^*i*^ ∈ [0, 1], with *A*^*i*^ = 0 corresponding to ECCs, *A*^*i*^ = 1 to MCCs and the intermediate values values represents different phenotypic states on the EM spectrum.

## 3. The macroscopic equations

Having established the individual–based description of cancer-cell dynamics, we next ask whether a macroscopic formulation can be obtained to capture the spatio-temporal evolution of the heterogeneous tumour mass as a whole. To this end, we adopt the kinetic–to–continuum methodology developed in [26, 8, 5, 4, 46]: first a kinetic equation is formulated, and then its macroscopic limits are derived, yielding the corresponding PDEs. While the cited works emphasise the analytical justification of the macroscopic limit, our present focus is confined to the construction of the kinetic model and the formal passage to the continuum scale.

### 3.1. The kinetic formulation

The first step to that is to derive the kinetic equation of the microscopic system of equations where we have assumed that the total mass of the system is conserved. Since we are working with both ODEs and SDEs we derive the kinetic equation of the number/probability density function *f* = *f* (*t, x, v, a*) : ℝ × ℝ^2*d*^ × [0, 1] → ℝ, according to the classical theory established for the Fokker-Planck or Vlasov equation [26, 61, 24, 21]. Let us note that the rigorous derivation of the so called *mean-field* equation has been studied for similar systems without the inclusion of the phenotypic variable and here, we give an overview of the steps needed for the derivation, inspired by the works of [2, 10, 11].

We assume that the system of *N* interacting cells of (2.1)-(2.3) is endowed with independent and commonly distributed initial data 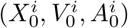 with 1 ≤ *i* ≤ *N* . Hence, due to the symmetry of the initial configuration and the evolution, all cells have the same distribution on ℝ^2*d*^ × [0, 1] at time t, which will be denoted as 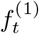. The cells get correlated due to the non-local term in (2.2), but since the pairwise action of two cells *i* and *j* is of order 1*/N*, it seems reasonable that two of these interacting particles become less and less correlated as *N* → ∞. This phenomenon is called propagation of chaos [51]. Thus, the *N* interacting stochastic process (*X*^*i*^, *V* ^*i*^, *A*^*i*^)_*t≥*0_ will behave approximately the same as the process 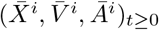 as *N* → ∞:

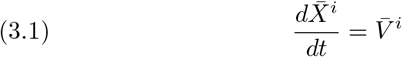

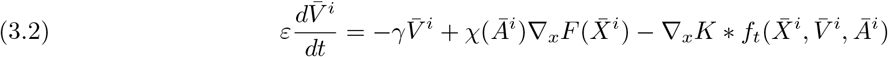

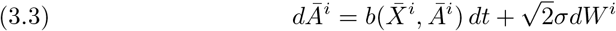

where *f*_*t*_ is the related probability density function. The initial datum of the system (3.1)-(3.3) is the same as the original one, so the stochastic processes 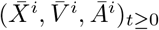 are independent from each other and identically distributed. Henceforth, from Itô’s formula we can derive the kinetic equation (see e.g. [4, 46]) for the evolution of the common probability distribution *f* that reads as follows:

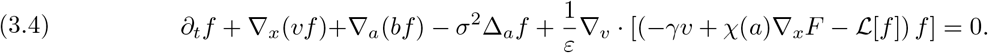

Here, with ∇_*x*_, ∇_*v*_, ∇_*a*_ we denote the gradient with respect to the position, velocity and phenotypic field, respectively, and with Δ_*a*_ the Laplacian operator in the phenotypic space. The operator [*f*] is formulated as follows:

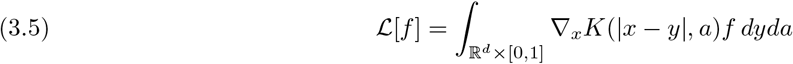

The existence of solutions of the kinetic equation needs more exploration for different choices of the functions *F, b* and the kernel *K*, but as long as a solution exists then the empirical measure, defined as

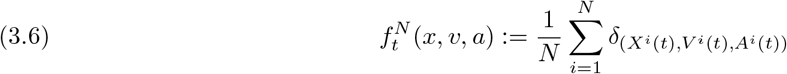

satisfies the kinetic equation in the distributional sense, i.e. 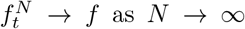. We provide more information on how the empirical measure satisfies (3.4) in Appendix A.

### 3.2. Derivation of the macroscopic equations

The next step now is to derive the macroscopic equations starting from the kinetic level. Following the works in [4, 5], we define the following ansatz for the probability density function *f* as:

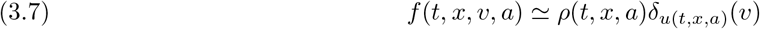

where *ρ* : ℝ × ℝ^*d*^ × [0, 1] → ℝ and *u* : ℝ × ℝ^*d*^ × [0, 1] → ℝ^*d*^ are the macroscopic density and the mean velocity of the particle system with

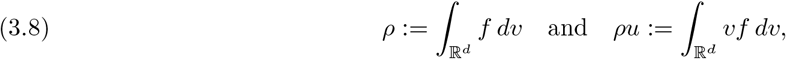

which are the first and second moments of *f* in respect of the velocity field. Taking now the integral over the velocity *v* in (3.4) and using (3.7) we have:

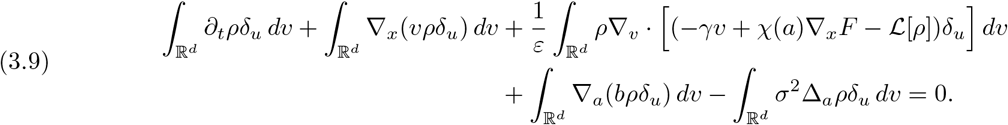

and using the properties of the Dirac function, for *ε >* 0, we arrive at

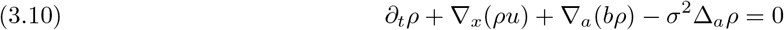

which is the macroscopic equation for the density of cancer cells. Similarly now we multiply (3.4) with the velocity *v* and take the integral over the velocity field.

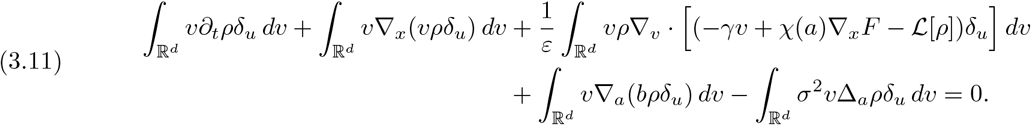

which gives, for *ε >* 0,

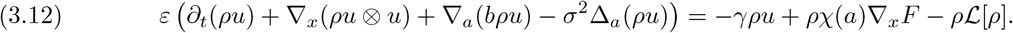

This equation describes the evolution of the momentum *ρu*. The overall system of macroscopic equations (3.10) and (3.12) derived, resembles an Euler-like system in space and time. The difference comes with the addition of the phenotypic space transitions that describe the continuous change between the epithelial and mesenchymal character of the cancer cell density *ρ* along side the momentum *ρu*. Note that inertia coefficient *ε* play a crucial role in the momentum equation (3.12). In this work we will study two separate regimes where *ε* = 1 and *ε* → 0.

In the inertia-less regime, i.e. *ε* → 0, the system (3.10)-(3.12) becomes:

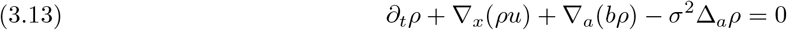

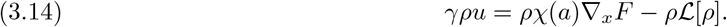

It is easy now to combine equations (3.13) and (3.14) and arrive at a single equation for the macroscopic density that is

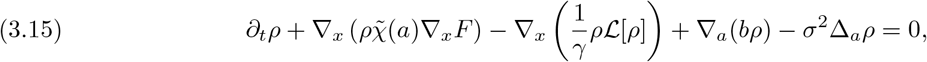

where 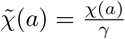. Equation (3.15) is an aggregation/diffusion advection equation for the density *ρ*(*t, x, α*) of the cancer cells. Notice that once more, as in (3.10)-(3.12), we have advection and diffusion-like terms for both the phenotypic space, and the physical space dimension. The second term of (3.15) describes the directed movement of cells towards higher values of the external field *F*, which can be interpreted as chemotaxis or haptotaxis in the context of cancer invasion. The third term represents the diffusion and aggregation of cancer cells due to their non-local interactions with their neighbouring cells. The last two terms, as before, depict the phenotypic transitions of cancer cells through both the field *b* or by simple diffusion.

### 3.3. Calculation summary

From the derivation presented in the previous section, we focus only on two different regimes for the coefficient *ε*, for the spatio-temporal evolution of cancer cell populations. The first one, for *ε* = 1, is an Euler-like system of equations that reads:

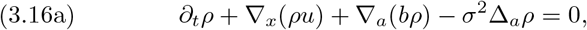

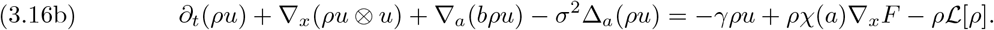

This formulation is based on the assumption that cancer cells can be represented as a complex fluid-like medium, with their behaviour regulated by their corresponding phenotypic state.

In addition to (3.16a)-(3.16b), we derived a single advection–aggregation–diffusion equation (AADE), when *ε* → 0, that describes the dynamics of the tumour cell density while explicitly accounting for the phenotypic heterogeneity of the cell population; this equation reads:

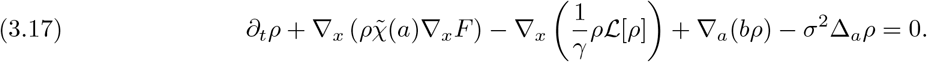

This second model is aligned with classical phenomenological approaches commonly found in the mathematical oncology literature [9, 15, 57, 42, 40, 13].

The aim now, is to investigate these two modelling approaches, highlight their differences, and assess their respective strengths and limitations in capturing the heterogeneous nature of cancer cell populations. In particular, we aim to explore their capacity in representing phenotypic plasticity and transitions mediated by EMT and MET processes. Moreover, we will also numerically investigate the agreement between to original IBM model with its corresponding macroscopic description.

### 3.4. Cell interactions

For this modelling approach, it is necessary to define the form of the non-local interactions between cancer cells, as well as the influence of extracellular cues. As cancer cells with a pronounced epithelial phenotype exhibit stronger adhesion forces with neighbouring cells, whereas cells with a dominant mesenchymal phenotype primarily migrate and invade the surrounding tissue, the phenotypic transitions play a central role in these definitions. This distinction reflects biological observations, where mesenchymal-like cancer cells display enhanced motility compared to their epithelial counterparts. Specifically, regarding the formulation of the interaction potential, we adopt a decomposition into repulsive and adhesive components. A typical choice is the Morse potential [5], which, in this context, takes the form:

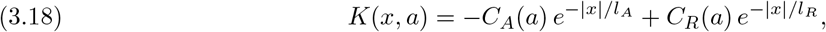

where *C*_*A*_, *C*_*R*_ *>* 0 and *l*_*A*_, *l*_*R*_ *>* 0 denote the adhesion and repulsion strengths and their corresponding interaction range/length. Here, the adhesion strength *C*_*A*_ is assumed to depend on the phenotypic state *a*, while the repulsion strength *C*_*R*_ is taken as phenotype-independent. The interaction range for adhesion is assumed to satisfy *l*_*A*_ *> l*_*R*_, consistent with the biological remark that ECCs are more capable of forming stable adhesive bonds, whereas this capability diminishes, if not completely eliminated, for MCCs.

Regarding repulsion, we introduce the assumption that all cancer cells exert equal strength repulsive forces, independent of their phenotypic state. Additionally, we consider the simplifying assumption that the repulsion forces act at the cell centres, which allows the repulsive term of the kernel to be approximated by a Dirac mass. Under these assumptions, the interaction potential is revised as:

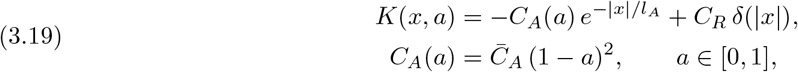

where 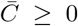 constant, and where the value *a* = 0 corresponds to ECCs and *a* = 1 to MCCs. The specific choice for the functional form of *C*_*A*_(*a*) can be different, provided that it reflects enhanced adhesive capabilities of ECCs relative to MCCs.

In addition, we specify the form of the extracellular signal *F* that governs the directed migration of cancer cells. The numerical simulations presented in the next section focus on haptotactic movement, where cells migrate towards regions of higher ECM densities. The ECM is assumed to remain stationary, i.e. immovable, and to be degraded by the cancer cells. We hence introduce the ECM density via the following equation:

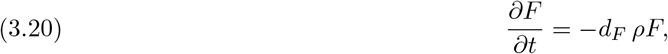

where *d*_*F*_ ≥ 0 represents the rate of ECM degradation induced by the cancer cells. Note that, we assume the degradation rate to be the same among also phenotypes. This is a simplification, since we expect cells with enhanced mesenchymal-like characteristics to degrade the ECM faster, in order to facilitate their motion. The directed motion of the cells, driven by the gradient of the ECM, is regulated by the haptotactic coefficient *χ* = *χ*(*a*) : [0, 1] → ℝ, which is dependent on the phenotypic state *a* ∈ [0, 1]. This dependency is introduced to capture the enhanced migratory capability of MCCs compared to ECCs. The functional form of *χ* is chosen as:

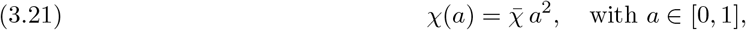

with 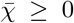 is constant, reflecting the quadratic increase of migratory potential with the mesenchymal phenotype.

The model further incorporates phenotypic transitions among cancer cells, governed by the function *b*, see (3.16a) and (3.17). The processes of EMT and MET are influenced by the local concentration of TGF-β, with EMT being promoted in regions where TGF-β is sufficiently abundant, while MET occurs in regions where its concentration falls below a given threshold. The density of TGF-β is denoted by *m* = *m*(*t*, **x**) : ℝ × ℝ^*d*^ → ℝ, which diffuses through the extracellular space according to:

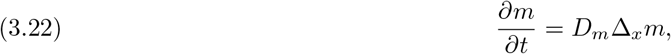

where *D*_*m*_ *>* 0 is the diffusion coefficient, assumed constant.

The dependence of the phenotypic transition coefficient *b* on the local TGF-β concentration is described through a drift function that dictates the transition rates for EMT and MET processes:

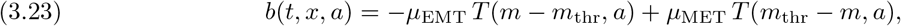

where *µ*_EMT_, *µ*_MET_ *>* 0 represent the respective transition rates, and *T* (*z, a*) : ℝ × [0, 1] → ℝ is the switching function that determines the availability of TGF-β for the corresponding phenotypic transition. The function *T* (*z, a*) is defined as:

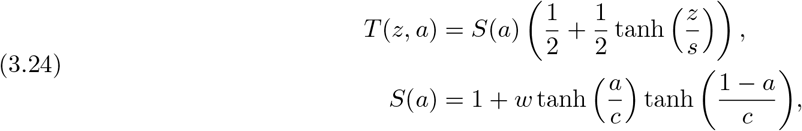

where the parameters *s, c* ≪ 1 control the sharpness of the switching behaviour and the modulation of the maximum transition speed, respectively. The function *S*(*a*) introduces a higher transition rate for intermediate phenotypic states, consistent with the modelling assumption that the dominant phenotypic states within the cancer cell population are the fully epithelial and fully mesenchymal types. This model approach provides a biologically motivated framework for the incorporation of phenotypic plasticity within the cancer invasion model.

## 4. Numerical experiments

The aim of this section is to gain further insight into the dynamical behaviour of the spatio-temporal phenotype-structured models given by (3.16a)-(3.16b) and (3.17), and to identify the potential phenomenological differences that may arise between these two modelling approaches.

To this end, we consider a series of numerical simulations designed to explore the behaviour of the proposed models under various experimental setups. These simulations focus on the interactions among ECC, hybrid epithelial–mesenchymal, and MCC populations, and on the role of phenotypic transitions in shaping the migratory patterns of the cells. It is important to note that the simulations presented here do not address the proliferation of the ECCs, as the models (3.17) and (3.16a)-(3.16b) are considered under the assumption of a fixed total cell population. Furthermore, we investigate the compatibility of the IBM (2.1)-(2.3) with the corresponding PDEs, to verify the validity of the limiting process.

The numerical solutions for the PDEs are obtained using an IMEX Finite Volumes/Finite Differences method, introduced in [36, 28, 38, 35]. In the case of the Euler-like model (3.16a)-(3.16b), the system includes the momentum term *ρu* and the phenotypic variable *a*, which is resolved using a finite difference discretisation scheme. For *a* ∈ [0, 1], the phenotypic grid points are defined as:

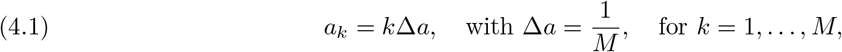

where *M* ∈ N denotes the total number of discretisation points in the phenotypic space. The discretisation in *a* leads to a system of spatio-temporal equations, which is then solved using the numerical scheme described in more detail in the Appendix B. For the numerical solution of the Euler-like system (3.16a)–(3.16b), we apply the schemes proposed in [53] adapted to the framework presented in [36].

The parameter values employed in the numerical experiments are summarised in Table 1. Unless explicitly stated otherwise, these parameters remain fixed across all simulations presented here. Moreover, since this is a comparative study between the two modelling approaches, all simulations are conducted using the same parameter sets. The spatio-temporal units of the parameters are not specified here, as the focus of the numerical study is on the qualitative features of the models rather than quantitative predictions, thus we work on dimensionless regime. All numerical simulations were performed using MATLAB R2025a [45].

**Table 1:**
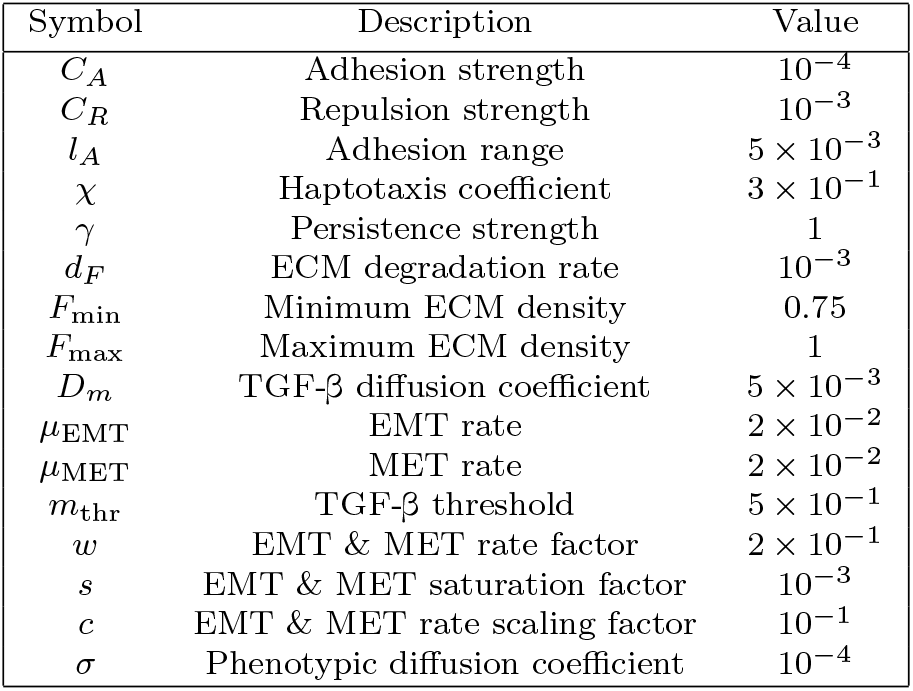
Parameter values used for the numerical simulations of the two models (3.16a)-(3.16b) and (3.17).

| Symbol | Description | Value |
| --- | --- | --- |
| $C_A$ | Adhesion strength | $10^{-4}$ |
| $C_R$ | Repulsion strength | $10^{-3}$ |
| $l_A$ | Adhesion range | $5 \times 10^{-3}$ |
| $\chi$ | Haptotaxis coefficient | $3 \times 10^{-1}$ |
| $\gamma$ | Persistence strength | 1 |
| $d_F$ | ECM degradation rate | $10^{-3}$ |
| $F_{\min}$ | Minimum ECM density | 0.75 |
| $F_{\max}$ | Maximum ECM density | 1 |
| $D_m$ | TGF-β diffusion coefficient | $5 \times 10^{-3}$ |
| $\mu_{\text{EMT}}$ | EMT rate | $2 \times 10^{-2}$ |
| $\mu_{\text{MET}}$ | MET rate | $2 \times 10^{-2}$ |
| $m_{\text{thr}}$ | TGF-β threshold | $5 \times 10^{-1}$ |
| $w$ | EMT & MET rate factor | $2 \times 10^{-1}$ |
| $s$ | EMT & MET saturation factor | $10^{-3}$ |
| $c$ | EMT & MET rate scaling factor | $10^{-1}$ |
| $\sigma$ | Phenotypic diffusion coefficient | $10^{-4}$ |

The numerical simulations are performed over a 2D spatial domain Ω = [−2, 2]^2^. For all phenotypic states, the cancer cell density *ρ* is assumed to be confined within this domain with zero Neumann boundary conditions. Zero-flux boundary conditions are also imposed at the boundaries of the phenotypic space. In Experiment 1, we adopt a coarse discretisation of the phenotypic domain with *M* = 10, reflecting the expectation of limited phenotypic diversity in the intermediate stages [47]. However, in Experiments 2 & 3, we use finer discretisations.

### Experiment 1 – Balanced EMT & MET haptotaxis flow

In this first experiment, we investigate the dynamics of the two derived PDE model models (3.17) and (3.16a)-(3.16b). In particular, in this scenario we aim to exhibit the interplay of EMT & MET and how these processes, affect the migratory patterns of the phenotype-dependent cancer cell population.

At the initial time, we consider a population comprised exclusively of cancer cells with a purely epithelial phenotype *a* = 0. The initial cell density is prescribed, for all **x** ∈ Ω, as:

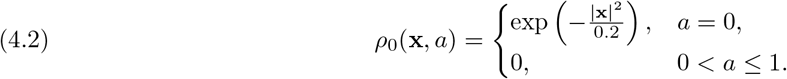

The initial condition for the ECM density is set such as, that there is a clear gradient structure oriented towards the bottom-right corner of the domain Ω. Namely, we set

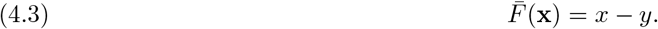

Subsequently, the ECM density values are normalised

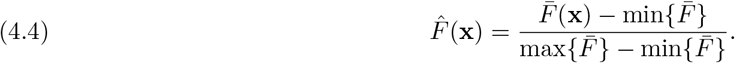

and, lastly, are brought within a biologically realistic range [*F*_min_, *F*_max_]

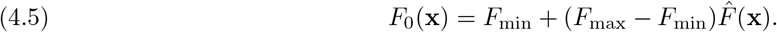

The range of ECM density we use in all experiments is [*F*_min_, *F*_max_] = [0.8, 1.25].

The initial TGF-β distribution is generated randomly across the domain, using a construction process originally presented [22] and given in more detail in Appendix C and Figure 9. The TGF-β threshold, *m*_thr_ = 0.5, is chosen such that it is within the range of the initial TGF-β configuration. This will lead to a balanced behaviour of the EMT & MET processes.

Furthermore, we assumed that all cancer cells, regardless of their phenotype, degrade the ECM at the same rate and that the TGF-β concentration diffuses with time.

Under this spatially heterogeneous TGF-β distribution, we observe that EMT is initially triggered only on the right-hand side of the primary tumour, where the TGF-β concentration exceeds the transition threshold. This leads to the progressive generation of cancer cell populations spanning the full phenotypic spectrum. At the intermediate time *t* = 20, both models once again capture the onset of MCC migration towards the bottom-right region of the domain, consistent with the ECM gradient. However, due to the diffusion of TGF-β, the migrating MCCs begin to encounter areas where the concentration falls below the EMT threshold. This shift promotes the activation of the reverse transition process, namely MET.

The effects of MET become clearly apparent at the final time *t* = 40. In particular, we observe an increase in the concentration of partial epithelial–mesenchymal cells in the regions previously invaded by the MCCs. Moreover, the reversion to an epithelial-like phenotype in these areas initiates the formation of a new, smaller tumour colony composed of ECCs. It is worth noting that, both models demonstrate directed movement along the ECM gradient, preserving their migratory behaviour.

Comparing the outcomes of the two models, no major qualitative differences in the migration patterns are observed. The spatial patterns of migration and phenotypic transition are consistent across the two frameworks, and no substantial differences in the resulting dynamics can be discerned from the simulations presented in Figures 2 and 3. This observation is likely dependent on the initial conditions and the parameter setup chosen in this Experiment. However, for the rest of our numerical tests, we will focus only on the AADE model (3.17), due to the computational complexity and runtime that the Euler-like system requires.

**Fig. 2:**
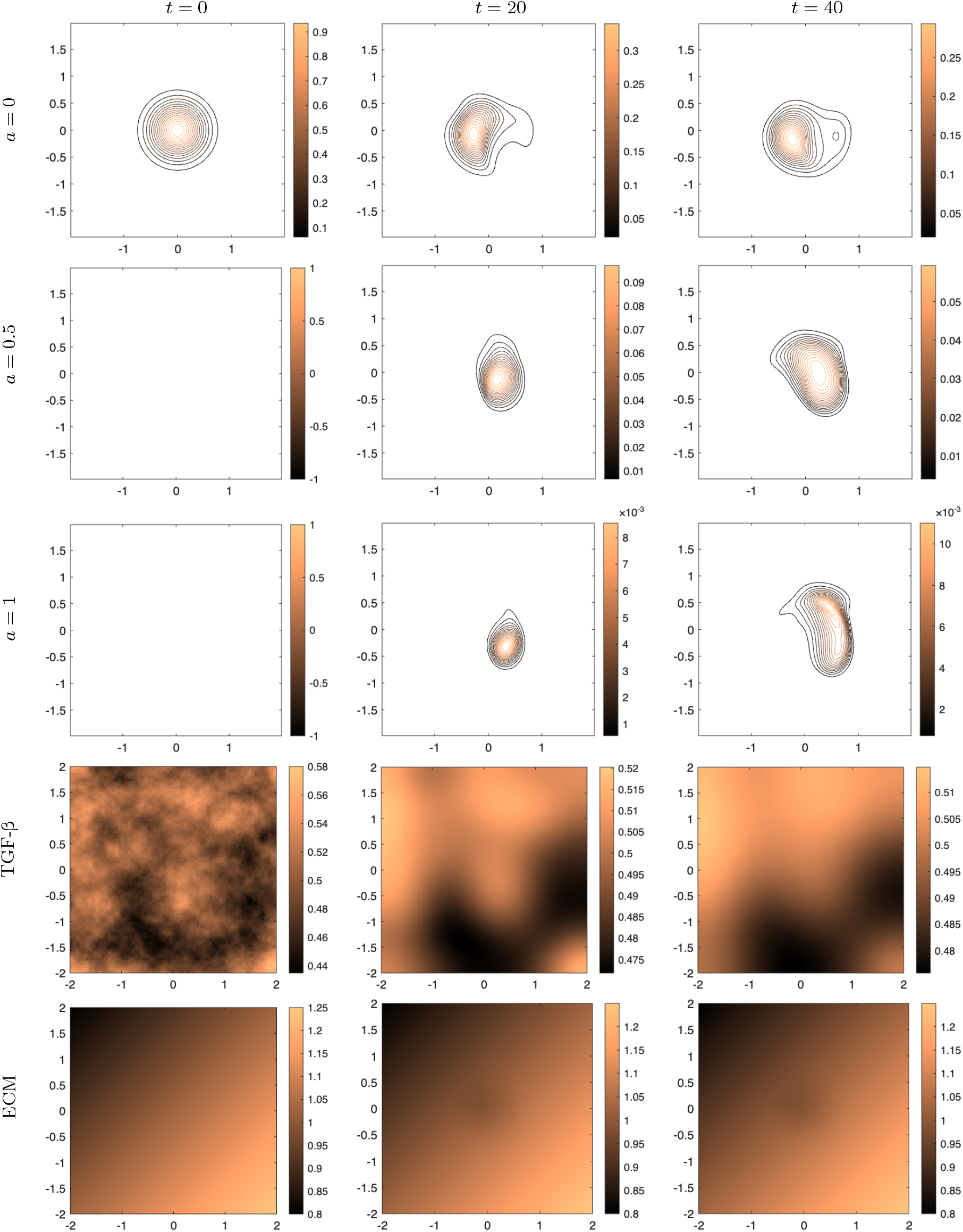
Experiment 1 — Haptotaxis flow in balanced TGF-*β* concentration: plots of the AADE model (3.17). EMT occurs only in selected regions, resulting in small MCC (*a* = 1) colonies that progressively migrate, due to haptotaxis, towards the bottom corner of the domain Ω. At *t* = 40, MET becomes more evident, leading to the formation of a smaller ECC (*a* = 0) colony adjacent to the original one. The qualitative behaviour is consistent with that of the Euler-like system shown in Figure 3.

**Fig. 3:**
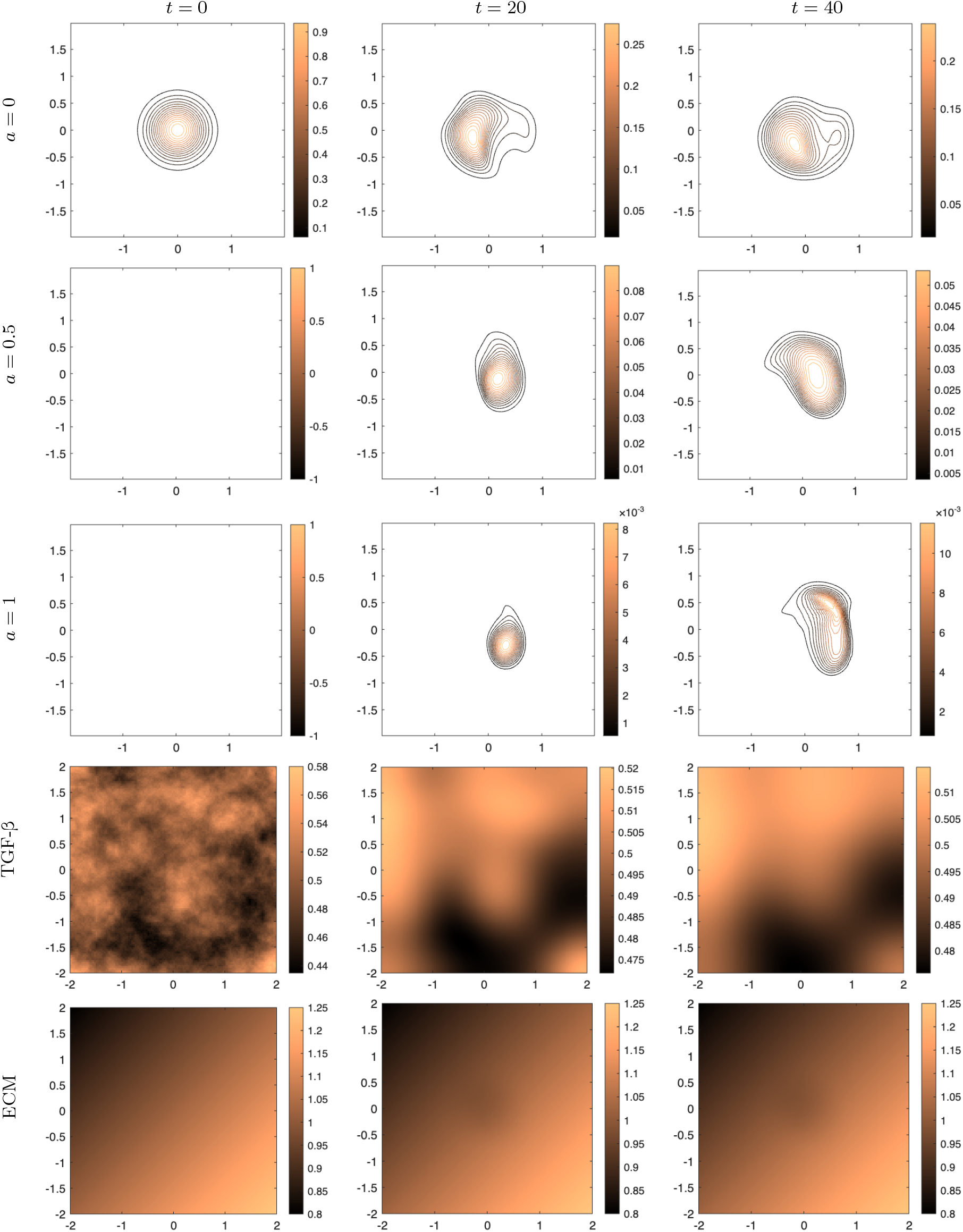
Experiment 1 — Haptotaxis flow in balanced TGF-*β* concentration: plots of the Euler-like system (3.16a)–(3.16b). EMT occurs only in selected regions, resulting in small MCC (*a* = 1) colonies that progressively migrate towards the bottom corner of the domain Ω. At *t* = 40, MET becomes more evident, leading to the formation of a smaller ECC (*a* = 0) colony adjacent to the original one. The qualitative behaviour is consistent with that of the AADE model shown in Figure 2.

### Experiment 2 – Haptotaxis flow towards a peak

In the second numerical experiment, we modify the initial configuration of the ECM to introduce a spatial structure resembling a “mountain range” with a gradually increasing ridge along the peak. The initial condition for the ECM density is shown in Figure 4 and is given by:

**Fig. 4:**
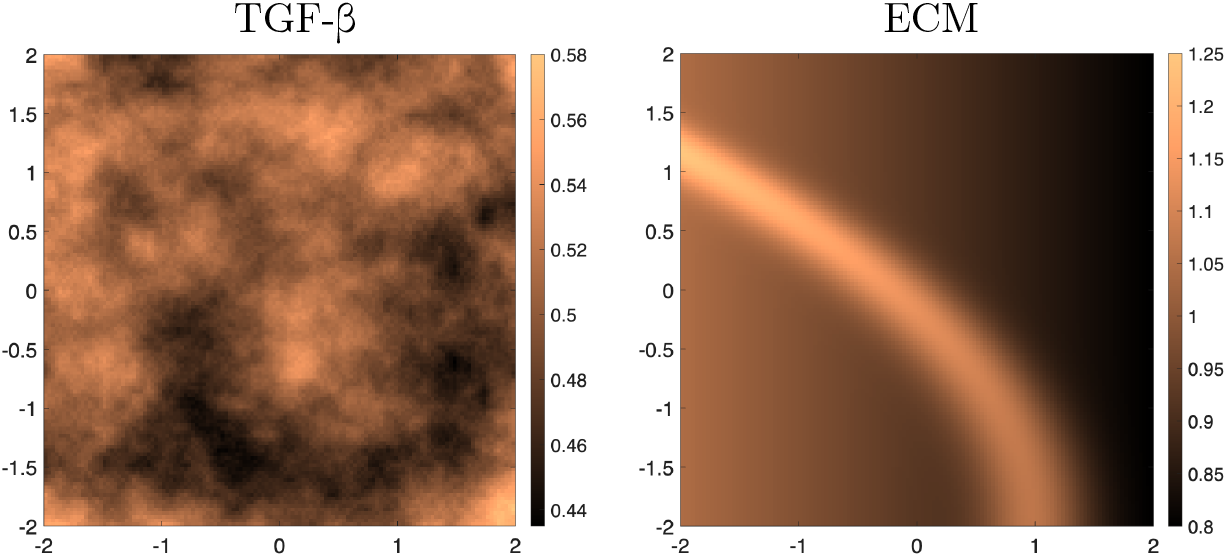
Experiment 2 — Haptotaxis flow towards a peak: plots of the initial density distributions of TGF-β and ECM, used in the results shown in Figure 5. The ECM density resembles a “mountain” range with a gradually increasing ridge along the peak.

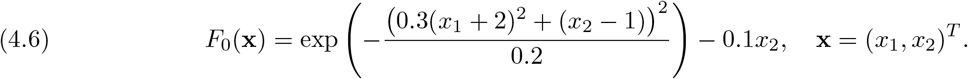

For the density of TGF-β we use the same initial distribution as was done in Experiment 1. The initial condition of the TGF-β concentration is shown in Figure 4.

As in Experiment 1, the initial cancer cell population is composed exclusively of ECCs. However, in this case, the population is spatially positioned at the “base” of the ECM ridge, located near the top-right corner of the domain Ω. The initial condition for the cell density is prescribed as follows, for **x** = (*x*_1_, *x*_2_) ∈ Ω:

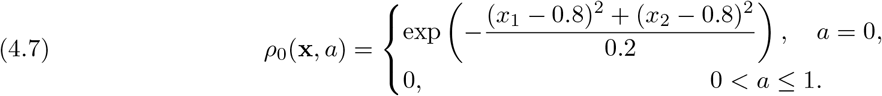

In addition to the change of the extracellular environment, in this experiment we aim to compare the AADE model (3.17), with its corresponding IBM description (2.1)-(2.3) where *ε* → 0. For the simulation of the IBM, we set the initial population of ECCs *N* = 3 × 10^3^ which is normally randomly distributed, again at “base” of the ridge, as we did for the macroscopic model. For the numerical solution of the IBM, we use a second order Runge-Kutta method to solve the ODE-part of the IBM, and an Euler-Maruyama scheme for the SDE-part of the model, in order to match the accuracy of the numerical method used for the PDE model. Let us note, that the IBM draws information from the PDE formulation of the ECM (3.20) and TGF-β (3.22) densities. Thus, for the numerical solution of this hybrid system of equations we use similar techniques as in [32].

In Figure 5, we showcase the comparative results of the spatio-temporal evolution, between the AADE and its corresponding IBM model. In particular, each column corresponds to a different range of phenotypic states, starting from cells that adopt phenotypes closer to the epithelial side of the spectrum (*α* ∈ [0, 0.25]), moving to partial epithelial-mesenchymal like cells (*α* ∈ (0.25, 0.5] and *α* ∈ (0.5, 0.75]) and finally, phenotypes closer to mesenchymal side of the spectrum (*α* ∈ (0.75, 1]).

**Fig. 5:**
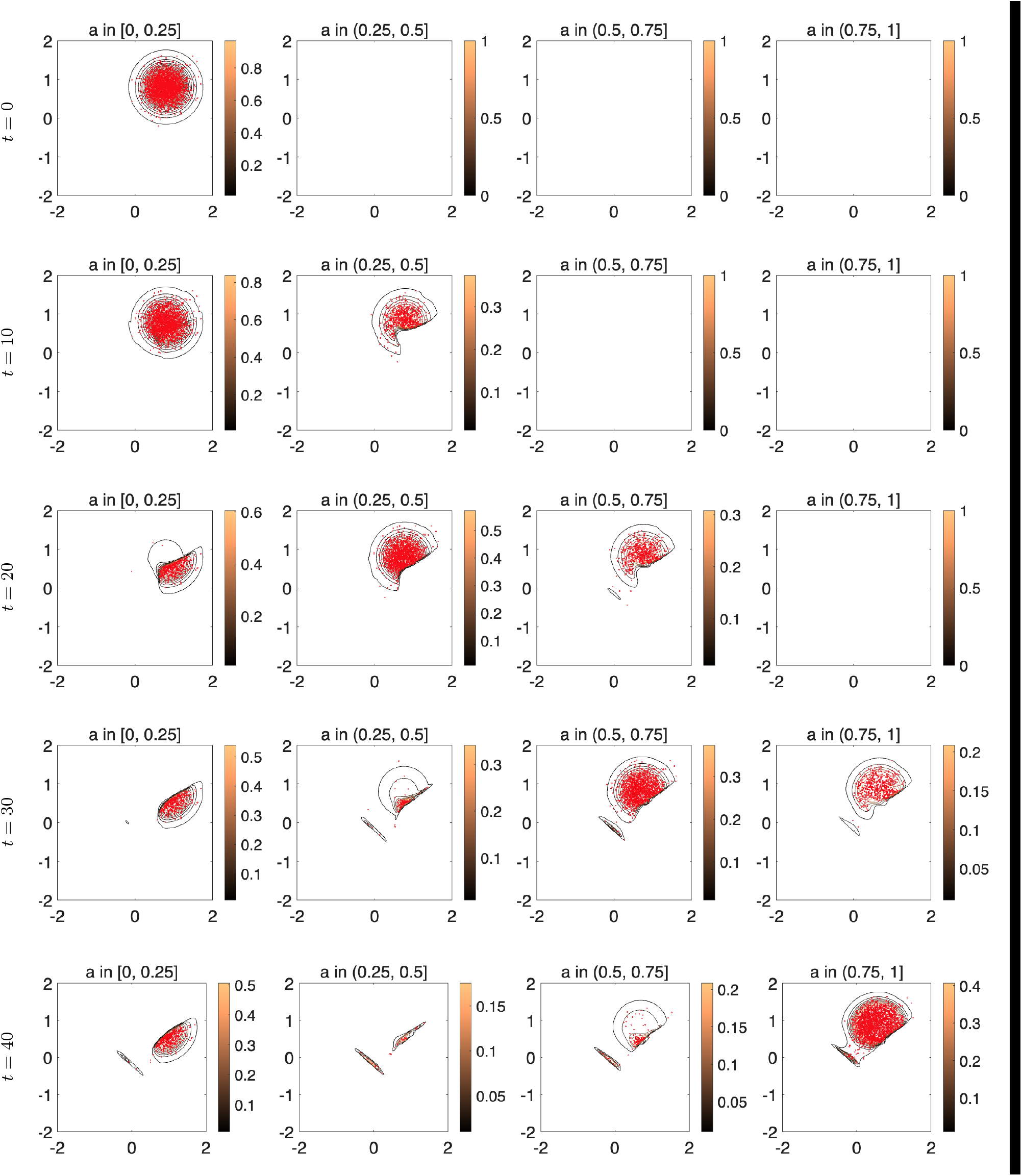
Experiment 2 — Haptotaxis flow towards a peak: plots of the AADE model (3.17), in contrast to the IBM (2.1)-(2.3). Comparison between the IBM, shown by red markers, and the macroscopic AADE density, shown by isolines, for four phenotypic bins. EMT and MET lead to the formation of two separate tumour colonies. The first is associated with the initial ECC population (*a* = 0), from which newly formed cells, with enhanced mesenchymal characteristics, migrate towards the top-left of the domain Ω due to the self-generated ECM gradient. The second colony results from escaping MCCs reaching the peak of the “mountain” in the centre of the domain Ω.

We observe that the numerical results of the IBM, qualitatively match the density evolution of the AADE model, which in effect verifies the validity of the limiting process performed in Section 3, from the microscopic to the macroscopic level.

The numerical results obtained from both the AADE model and IBM reveal two distinct invasion dynamics. Firstly, we observe the progressive emergence of a cell population, of enhanced mesenchymal-like characteristics, i.e. *α* ∈ [0.5, 1], that detaches from the original tumour mass and migrates actively towards the peak of the ECM ridge. Upon reaching the nearest local maximum of the ECM, these mesenchymal-like cells continue to migrate along the ridge, progressing towards the top-left region of the domain. This invasion process facilitates the formation of a secondary tumour colony, driven by both active mesenchymal migration and the phenotypic reversion promoted by low TGF-β concentrations in the central region of Ω. The occurrence of MET in these regions supports the generation of new epithelial-like cells, i.e. *α* ∈ [0, 0.5], thus contributing to the development of a mixed phenotypic structure at the metastatic site. These dynamics are captured in both models at time *t* = 40, as illustrated in Figure 5, where two spatially distinct tumour populations—the primary and the metastatic—are clearly visible.

The second mechanism of invasion is observed within the original tumour region. The epithelial-like population is initially located in a region of low ECM density, lacking a well-defined gradient for directed migration. However, due to ECM degradation by the cancer cells, we observe the emergence of a self-generated ECM gradient. This gradient subsequently enables cell motility, leading to movement of primarily mesenchymal-like cells, towards the top-left corner of the domain. The formation of this self-generated pathway is particularly evident in Figure 5 at time *t* = 40.

### Experiment 3 – Microscopic versus macroscopic models and phenotypic plasticity

In this experiment, we investigate two specific questions: whether the IBM (2.1)-(2.3) and the corresponding AADE (3.17) produce the same spatial and phenotypic distribution, similarly to Experiment 2, and how the sharpness of the TGF-β-dependent EMT switch function (3.24) affects the distribution of cancer cells across the EM spectrum.

To this end, we perform two sets of simulations, one with a steep phenotypic function and one with a smooth (3.24). The parameter, *s*, that controls the steepness of the switch function, is chosen in the first case to be sufficiently small so that the phenotypic response is almost instantaneous when the TGF-β threshold value *m*_thr_ is reached. In the second case, a larger value of *s* leads to a more gradual response to TGF-β. The TGF-β distribution that was used in both experiments and is shown, along with the two switch functions, in Fig. 6, where now the TGF-β threshold is chosen as *m*_thr_ = 0.9.

**Fig. 6:**
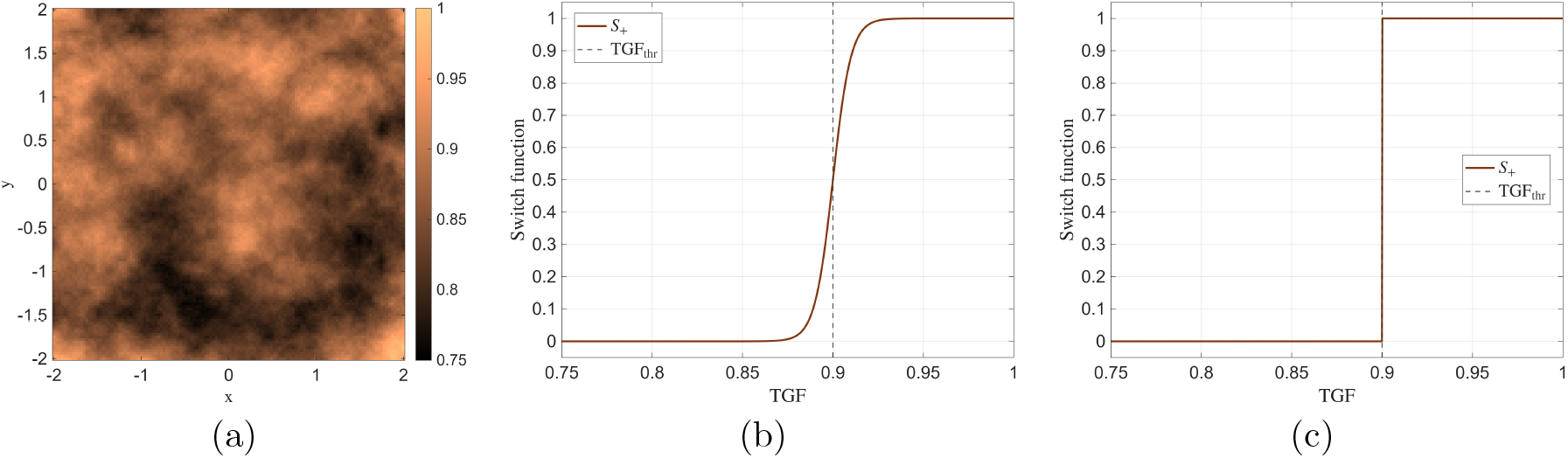
Experiment 3 — TGF-β distribution and TGF-β-dependent switch functions. (a) The fixed spatial distribution of TGF-β used in both the sharp and smooth switch simulations. The vertical dashed line indicates the threshold value *m*_thr_. (b) Smooth switch function. (c) Sharp switch function.

For the sake of clarity of comparison between the two models, we assume a simplified version of the ECM density model (3.20), where the cells do not degrade the matrix, implying that the ECM density is stationary with *F* (*t, x*) = *F*_0_(*x*). The migration of the cells is still affected by the gradient of the ECM through (2.2). The same holds true for the TGF-β density distribution, which is kept fixed throughout these simulations. For the initial conditions of the cancer cell density *ρ*, the ECM and the TGF-β, we use the same setting as in Experiment 1. Similarly, for the initial distribution of the IBM, we randomly place *N* = 10^4^ cells close to the centre of the domain, using a normal distribution N(*µ, σ*^2^) with mean *µ* = 0 and standard deviation 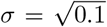. We also assume no cell-cell interactions, in order to focus only on the dynamics of phenotypic transitions. All remaining parameters are as in Table 1.

The simulation results in Fig. 7 and Fig. 8 show the comparison between the IBM and the macroscopic PDE model. The initial cancer cell population consists exclusively of epithelial-like cells, *a* = 0. The phenotypic variable remains continuous, *a* ∈ [0, 1], but for visualisation purposes we have, yet again, grouped both the individual cells and the AADE macroscopic density into four bins: *a* ∈ [0, 0.25], *a* ∈ (0.25, 0.5], *a* ∈ (0.5, 0.75], and *a* ∈ (0.75, 1]. The first and last bins correspond to cells with dominant epithelial- and, respectively, mesenchymal-like, character; the two middle bins represent partial EM phenotypes.

**Fig. 7:**
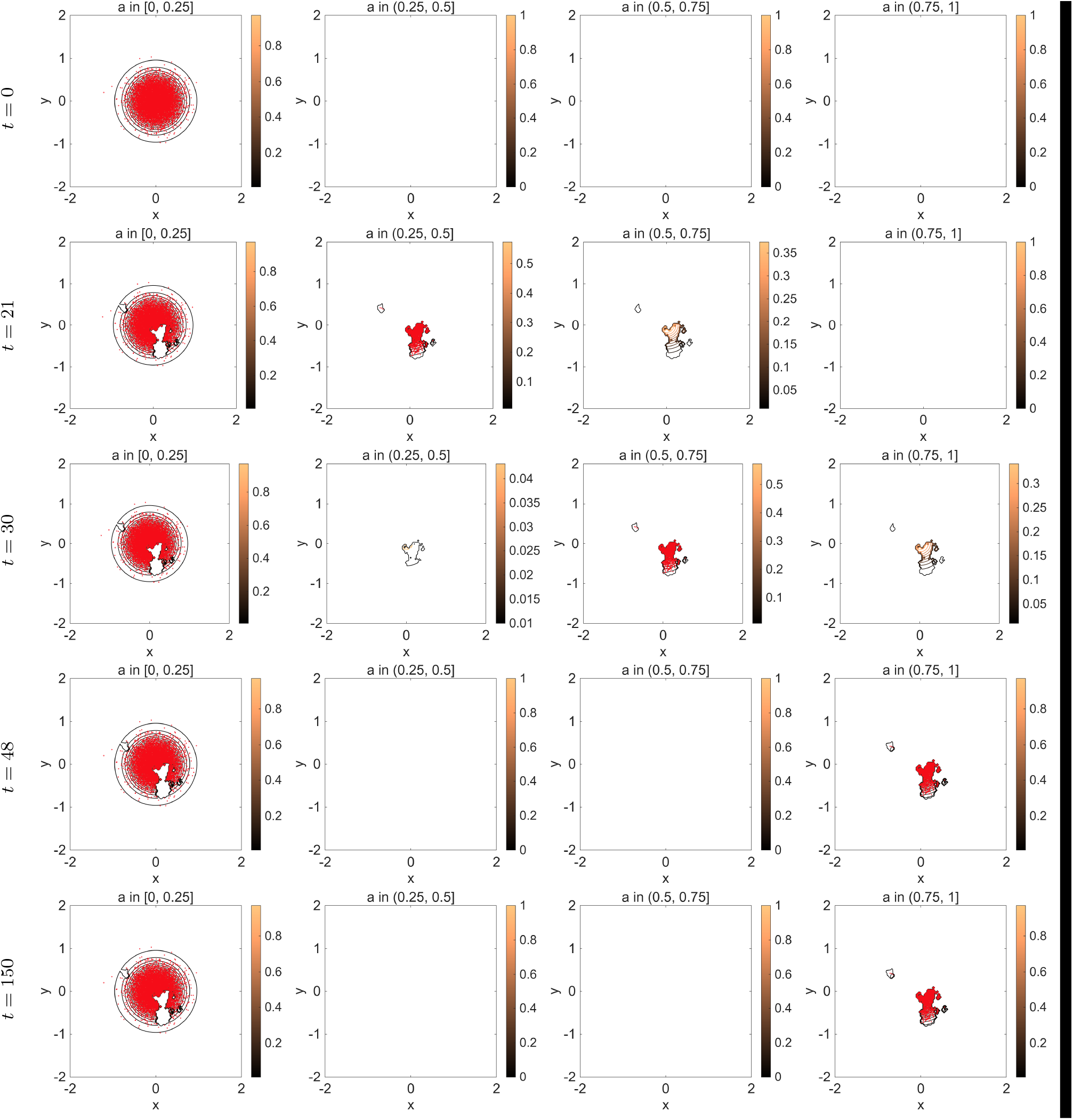
Experiment 3 — Microscopic vs Macroscopic under a sharp TGF-β-dependent switch. Comparison between the IBM, shown by red markers, and the macroscopic AADE density, shown by isolines, for four phenotypic bins. The sharp switch produces an almost binary EMT response, with cells concentrated predominantly in the epithelial-like and mesenchymal-like bins, while the intermediate bins remain almost empty.

**Fig. 8:**
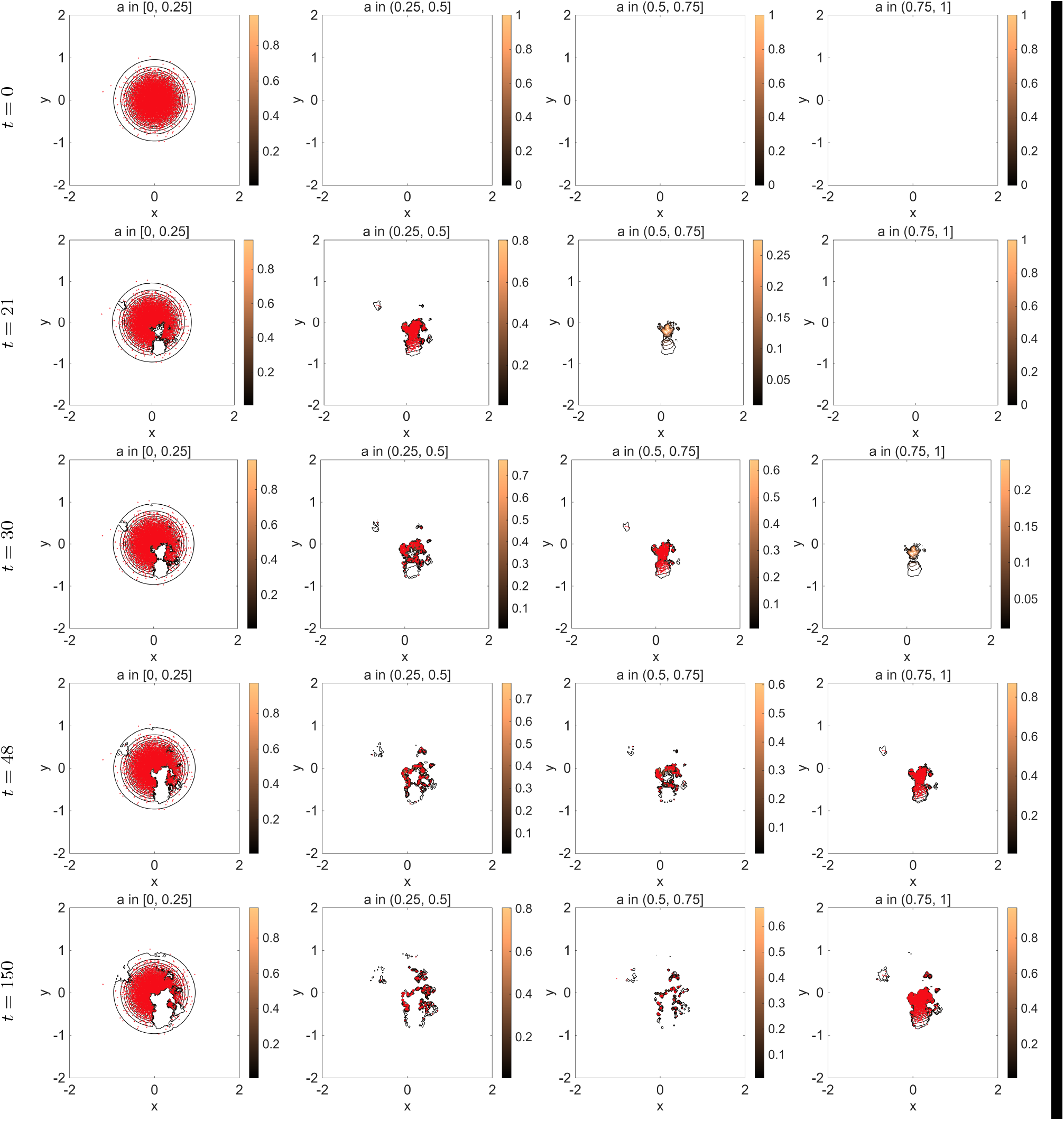
Experiment 3 — Microscopic vs Macroscopic under a smooth TGF-β-dependent switch. Comparison between the IBM, shown by red markers, and the macroscopic AADE density, shown by isolines, for four phenotypic bins. The smooth switch gives rise to a broader distribution of cells across the EM spectrum, including partial EM phenotypes.

**Fig. 9:**
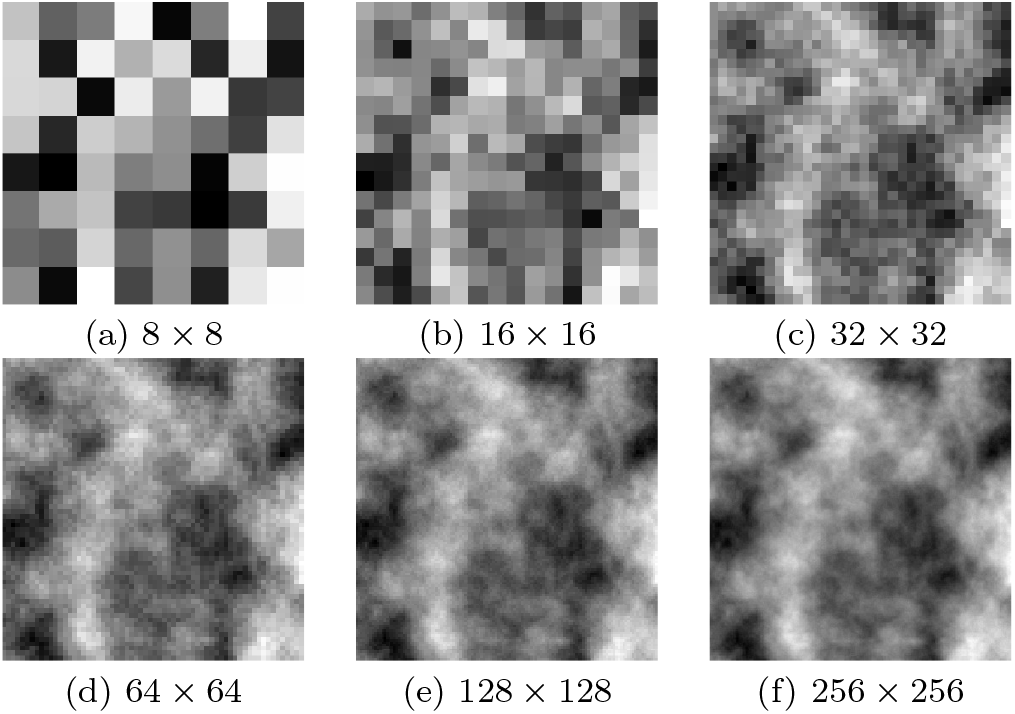
Construction of the (initial condition) TGF-β concentration employed in Experiments in Section 4. The process starts with random values over an 8 × 8 grid; these introduce the basic structure of the ECM. In every stage of the construction process, the grid is bisected with the new values attained by averaging the neighbouring values of the previous coarser grid with the addition of some additive noise. The process stops when the required grid size is reached.

The individual cells are represented by red markers, while the macroscopic density is represented by isolines. The agreement between the two descriptions is very strong in both simulations. At the initial time, both the individual cells and the PDE density are concentrated in the epithelial-like bin and occupy the same spatial region. As time evolves, the PDE isolines continue to follow closely the spatial distribution of the individual cells. In particular, the isolines correctly identify the location of the epithelial-like population, the emergence and migration of the more mesenchymal-like population, and the presence (or absence) of intermediate phenotypes.

The agreement is particularly clear in regions of non-negligible mass. In these regions, the isolines of the AADE model engulf the same cell clusters generated by the IBM, showing that the continuum density reproduces both the spatial localisation and the phenotypic composition of the underlying cell population. Small discrepancies appear only near low-densities or isolated cells, which is expected due to the finite cell number in the IBM. Overall, the agreement observed in Figs. 7 and 8 supports the compatibility of the IBM with the phenotype-structured AADE.

In the sharp-switch case, Fig. 7, the population rapidly separates into two dominant phenotypic families. A large part of the population remains in the more epithelial-like bin *a* ∈ [0, 0.25], while cells located in regions of sufficiently large TGF-β concentrations, transition towards the mesenchymal-like bin *a* ∈ (0.75, 1]. The two intermediate bins are only transiently in time populated, due their drift along the *a*-dimension, (2.3) and remain almost empty at later times. Thus, the sharp switch produces a nearly binary EMT response, where cells behave either as epithelial-like or mesenchymal-like. The epithelial-like cells remain concentrated close to the initial tumour region, whereas the mesenchymal-like cells migrate towards the lower-right region of the domain, in agreement with their stronger haptotactic response discussed in the previous experiments.

In the smoother-switch case, Figure 8, the cell population is distributed across the phenotypic space more broadly. At early times, populations are observed in the bins *a* ∈ (0.25, 0.5] ∪ (0.5, 0.75]. The smoother switch allows cells to remain in partial EM states over longer simulated times. At the final time, *t* = 150, the population is distributed across the full EM spectrum, with the epithelial-like cells remaining closer to the original tumour region and the more mesenchymal-like ones migrating further away.

Biologically, this experiment indicates that the cellular response to pathways of switching signals may determine whether EMT is observed as an binary change between EM phenotypes, or as a gradual progression through partial EM states. In the steep-switch case, cells respond to TGF-β in an almost threshold-dependent manner: below the threshold they maintain epithelial characteristics, while above it they acquire a mesenchymal-like phenotype. This is consistent with experimental systems that are understood through end-point EM phenotypes, for example epithelial-like breast cancer cell lines such as MCF7 or T47D, compared with more mesenchymal-like cell lines such as MDA-MB-231 or SUM159 [1, 29]. In contrast, the smoother switch produces a gradual response to TGF-β, allowing cells to occupy intermediate EM phenotypes. This behaviour is closer to what has been reported in cancer cell lines such as H1975 and PMC42-LA, where partial EM phenotypes have been observed [59, 31, 30, 1]. This distinction is important, since partial EMT phenotypes have been associated with collective migration in cancer cells, and metastatic capacity. Henceforth, the shape of the switch function could be used as a clear adaptable criterion when modelling different cell lines, and the sharpness parameter could be the point of inference, when dealing with experimental data.

## 5. Discussion

The objective of this study was to explore the relationship between IBMs for cancer cell invasion and their corresponding macroscopic descriptions. In particular, we introduced an individual-based framework that incorporates several key biological mechanisms relevant to the invasion-metastasis cascade, including cell-cell and cell-ECM interactions, as well as phenotypic plasticity. The integration of phenotypic heterogeneity into a classical individual-based modelling approach serves to address the common limitation of phenotypic uniformity often encountered in the literature of off-lattice models, where individual cells are assumed to respond identically to environmental cues.

To obtain the corresponding macroscopic representations, we considered two distinct modelling approaches: one resulting in a single AADE, and another leading to an Euler-like system for the spatio-temporal evolution of the phenotype-dependent cancer cell population. Both modelling frameworks incorporate phenotypic transitions consistent with biological observations: mesenchymal-like cells exhibit enhanced migratory capabilities, while epithelial-like cells are largely non-motile but retain proliferative properties.

The first aim of the numerical simulations, presented in Section 4, was to demonstrate the diversity of spatio-temporal dynamics arising from the interaction of cancer cell phenotypes. A critical factor in this context is growth factor TGF-β, which acting as a trigger of EMT-MET, plays a central role in tumour progression and metastatic spread. In Experiments 1 & 2, the simulation results show that the proposed models are capable of reproducing the evolution of the full phenotypic spectrum-including ECCs, hybrid epithelial-mesenchymal cells, and MCCs—with each subpopulation exhibiting behaviour consistent with their corresponding biological expectations.

The second aim was to compare the two macroscopic formulations, namely the AADE model (3.17) and the Euler-like system (3.16a)–(3.16b). From a qualitative perspective, the results indicate no major discrepancies between the two. Both models successfully capture the underlying biological mechanisms and reproduce consistent macroscopic behaviour. Minor quantitative differences, such as those observed in Experiment 1, suggest future investigation—either in terms of model refinement or numerical resolution. Further studies are needed to clarify the conditions under which one modelling framework may be preferred over the other in capturing the invasion-metastasis cascade. One notable limitation, though, of the Euler-like model (3.16a)-(3.16b) is its higher computational cost, particularly in three-dimensional simulations. Identifying concrete scenarios where the predictions of the two models diverge meaningfully would provide valuable insight, informing the selection of modelling strategies for biological and clinical applications. In parallel, analytical investigations should be pursued, focusing on well-posedness, existence of solutions, and the identification of parameter regimes that lead to qualitatively different behaviours between the two modelling approaches. The integration of such analytical results with extensive computational simulations is essential for advancing our understanding of cancer invasion and metastasis.

Furthermore, the microscopic versus macroscopic Experiment 2 & 3 has provided a direct comparison of the continuum description at the level of the phenotype-structured cell population. In both the sharp and smooth switch simulations of Experiment 3, the individual cells and the AADE (3.17) of the same phenotype occupy the same physical space. This comparison supports both the derivation and the use of the AADE as a consistent large-number limit description of the underlying individual cell dynamics, including both spatial as well as phenotypic distribution of the populations.

This experiment also emphasises the biological relevance of the EMT switch function (3.24). When the response to the extracellular cue is steep/swift, the population separates into two disjoint families, with cells remaining at the extremes of the phenotypic space. This behaviour is consistent with cellular systems such as epithelial-like breast cancer cell lines MCF7 and T47D, as opposed to the more mesenchymal-like cell lines such as MDA-MB-231 and SUM159 [1, 29]. In contrast, the smoother response to the extracellular cues generates a broader EM distribution. This is closer to what has been observed in EMT models such as HMLE and MCF10A cells exposed to TGF-β, or in cancer cell lines such as H1975 and PMC42-LA, [59, 31, 1, 30].

These results support the interpretation of EMT as a form of EM plasticity, rather than a process that must always be reduced to a binary EM switch. In the present model, changing the sharpness of the TGF-β response is sufficient to move from an extreme endpoint EM regime to one in which partial EM phenotypes persist over time. This suggests that the regulatory response to TGF-β may be a contributing factor to the experimentally observed differences between cell systems with mainly epithelial and mesenchymal phenotypes and those displaying partial EM properties.

It is important to highlight that both macroscopic models were considered under the assumption of a fixed cell population, with no source terms representing proliferation. For applications involving experimental validation, the inclusion of proliferation mechanisms is necessary, especially given their dependence on phenotypic state. Moreover, the current modelling framework assumes continuous phenotypic transitions governed by the local concentration of TGF-β. In reality, phenotypic switching is not constant but occurs under favourable microenvironmental conditions. This limitation may be addressed either by reformulating the phenotypic advection term as a stochastic process, e.g. by considering two distinct thresholds for EMT and MET.

Also, the formal derivation of the limiting macroscopic PDEs should be investigated further. The monokinetic ansatz used to pass from the kinetic to the macroscopic level could be improved by identifying the fast and slow dynamics of the model, and through a Hilbert expansion derive macroscopic equations for the phenotype dependent cancer cell population, [18, 16, 41, 43, 60].

Finally, we remark that the IBM defined in equations (2.1)-(2.3) accounts only for phenotypic transitions along the epithelial–mesenchymal axis. However, the general framework allows for the inclusion of multiple biological traits by extending the phenotypic variable to a multidimensional space, *α* ∈ R^*n*^. For example, nutrient availability (e.g. oxygen concentration) could be incorporated alongside the epithelial–mesenchymal scale. Such an extension would enable the construction of richer macroscopic models capable of capturing the interplay between various biological factors and tumour progression. These directions represent a natural and biologically motivated extension of the present work.

## Appendix A. The empirical measure satisfies the kinetic equation

Using the definition of the empirical measure we can express (2.2) as follows:

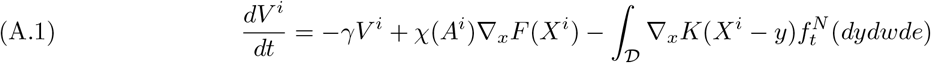

where *D* = ℝ^*d*^ × ℝ^*d*^ × [0, 1]. Now for any function *ϕ* ∈ C^1^(D) we have that:

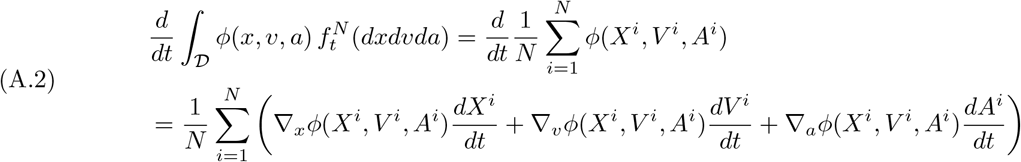

and if we use equations (2.1), (2.3), (A.1) to (A.2) using the definition (3.6) of the empirical measure we arrive:

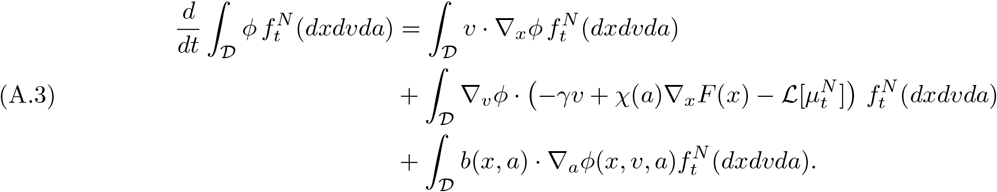

This yields that the empirical measure satisfies the kinetic equation (3.4) in the sense of distributions subject to the regularity of the functions *b, F* and the kernel *K*.

## Appendix B. Numerical solution of PDE systems

The advection-reaction-diffusion system in Section 4 is solved numerically using a second-order Implicit-Explicit Runge-Kutta Finite Differences, Finite Volumes (IMEX-RK FD-FV) numerical method. This method constitutes an extension of a previous method developed and employed in [36, 49, 35, 37, 38, 50, 22, 32], where we refer for most of the details. Here, we only discuss some of its components.

We consider, at first, a generic advection-reaction diffusion system of the form

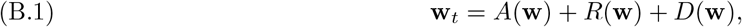

where **w** represents the analytical solution vector of the system, and *A, R*, and *D* are the advection, reaction and diffusion operators respectively. After spatial discretisations have taken place, we denote the corresponding semi-discrete approximation by **w**_*h*_, where the index *h* denotes the (maximal, if the space discretisation is non-uniform) spatial grid diameter. The semi-discrete solution **w**_*h*_ satisfies the following system of ODEs

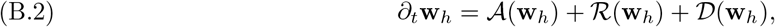

where the numerical operators *A, R* and *D* are (spatially) discrete approximations of the advection-reaction, diffusion operators, *A, R* and *D* in equation (B.1). Moreover, as the numerical scheme we employ is (partially) FV, raising its accuracy to the second order necessitates the use of flux limiters for the interface reconstruction of the numerical fluxes. Out of a large number of flux limiter options, we have found that the Minimized-Central (MC) limiter, see [55], constitutes a robust and efficient choice.

Let us note, that for the equations presented in Section 3.3, we first discretise the phenotypic domain and then move onto the spatial discretisation. In particular, we treat the phenotypic-advection, ∇_*a*_, and phenotypic-diffusion, Δ_*a*_, terms as reaction terms of the semi-discrete system (B.2).

Before solving (B.2), we re-organise its terms in implicit and explicit (IMEX splitting) and, accordingly, (B.2) takes the form

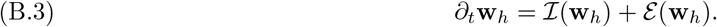

The actual IMEX splitting depends on the problem at hand, but in a typical case the advection *A* terms are treated explicitly in time, the diffusion terms *D* implicitly, and the reaction *R* terms either explicitly or implicitly, depending on their stiffness. In the problems that we encounter in this thesis, all reaction terms have been resolved explicitly in time.

The semi-discrete problem (B.3) is now solved using a diagonally implicit RK method for the implicit part *I*(*w*_*h*_), and an explicit RK for the explicit part *E*(**w**_*h*_). Altogether, we solve (B.3) using the additive RK scheme

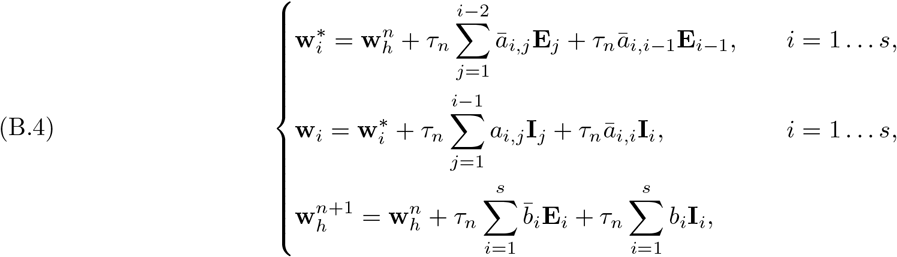

where *s* are the stages of the IMEX-RK method, **E**_*i*_ = **E**(**w**_*i*_), **I**_*i*_ = **I**(**w**_*i*_), *i* = 1 … *s*, and 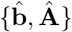, {**b, A**} are the coefficients for the explicit and implicit part of the scheme respectively. We refer the reader for the values of these coefficients to [34], where the Butcher’s Tableau of both second and third order methods are given. As a final stage of this method, we solve the linear system in (B.4) using the Iterative Biconjugate Gradient Stabilised Krylov subspace method, see [39, 54].

## Appendix C. Initial condition for numerical experiments

We give here a short description on the construction of the initial condition of the TGF-β concentration. A visual representation is shown in Figure 9, but the construction process goes as follows and more details can be found in [22, 32]. To begin with, an 8 × 8 matrix is created with entries taken from a standard normal distribution, *N* (0, 1). A number of refinement steps are taken until the resolution of the matrix reaches the desired (computational) resolution of the domain. At each stage, the size of the matrix is doubled to increase the resolution of the ECM. The entries to the new larger matrix are obtained from interpolating the values of the previous smaller matrix with the addition of a small amount of Gaussian noise. Therefore, as the ECM is refined it preserves the initial randomly chosen structure observed in the 8 × 8 matrix, with areas of higher or lower densities appearing in the same regions of the grid no matter what resolution is used. This procedure is extended into three dimensions. However, due to the increased computational time that three-dimensional simulations impose, a maximum refinement of 64 × 64 × 64 will be used for the ECM in this experiment.

